# Epigenetic Resilience to Early-Life Maternal Loss in African Savanna Elephants

**DOI:** 10.64898/2026.04.03.716431

**Authors:** Daniella E. Chusyd, Steven N. Austad, Janine L. Brown, Lanos Chisaka, Kasi Kalande, Claudia Lalancette, Milda Milčiūtė, Lisa Olivier, Innocent N’gombwa, Jonathan Sinyinza, Eric T. Klopack

## Abstract

Early-life trauma in humans, including maternal loss, is strongly associated with increased risk of chronic diseases, reduced life expectancy, and accelerated biological aging, as measured through epigenetic modifications, such as DNA methylation. Whether similar patterns occur in other long-lived, socially complex non-primate species, however, remains unknown. Elephants share key life-history traits with humans, including longevity, strong social bonds, and advanced cognitive abilities. Yet wild elephant populations face significant anthropogenic and environmental pressures, including poaching, habitat loss, and human-wildlife conflict that can result in maternal mortality and subsequent calf orphaning. We examined whether orphaning of elephant calves was associated with accelerated DNA methylation age and distinct epigenetic signatures. Contrary to our hypothesis and patterns observed in other species, orphaned African savanna elephants exhibited a younger DNA methylation age than non-orphans, no accelerated aging, and only limited differential methylation at CpG sites. At the genome-wide level, chronological age was not associated with differential CpG methylation after correcting for multiple testing. One interpretation of these findings is that elephants may have evolved mechanisms that buffer against epigenetic instability following stressful events. Investigating these protective mechanisms in elephants could inform strategies to mitigate the long-term health impacts of early-life trauma in humans.

**Significance Statement:** In most mammals, early-life adversity, including maternal loss is associated with shorter lifespans and widespread epigenetic alterations. In contrast, we found that orphaned African savanna elephants exhibited a younger epigenetic age compared to non-orphans and showed only a weak distinct epigenetic signature. This unexpected pattern may reflect environmental influences, such as living under human care, or evolutionary adaptations that buffer against epigenetic instability. If the latter is confirmed, such mechanisms could confer resilience to the epigenetic consequences of early-life adversity.

## Introduction

Elephants have long fascinated humans, in part because their complex social and cognitive capabilities resemble our own. Similar to people, elephants are long-lived, living into their 70s^1-4^ and 80s^2^ under certain circumstances. They rely heavily on social bonds, advanced cognition, and memory, and display behaviors associated with “theory of mind”, including self-awareness,^5^ cooperation with one another,^6^ mourning-like behavior,^7^ empathy,^8^ and consolation.^9^ Furthermore, elephant mothers have been observed burying deceased calves.^10^ Because of these psychosocial similarities, elephants are an important comparative species with strong potential for investigating how early-life experiences, like maternal loss, influence later-life health outcomes.

Wild elephant populations are subject to intense anthropogenic pressures, most notably ivory poaching and human-elephant conflict arising from competition over shared resources. At the peak of the poaching crisis in the early 2000s, 25,000 elephants was estimated to be killed in a year.^11^ Although elephants have well-documented social structures and behavioral responses to loss that include grieving, mourning, and consolation, the scale and persistence of anthropogenic pressure can be overwhelming, leading to behaviors similar to post-traumatic stress disorder (PTSD) in people. In one example, adolescent orphaned male elephants in South Africa, lacking the presence of older adult males, were uncharacteristically violent, killing over 100 rhinoceroses (an aberrant behavior for elephants).^12^ In addition, male elephants with PTSD-like behaviors were responsible for 90% of all male elephant deaths in their community, compared with 6% in relatively unstressed communities.^12^ Given the importance of social bonds in elephant society, coupled with elephants’ long-term memory,^13-15^ it is reasonable to posit that orphaned calves would experience trauma.

Humans who experience adverse childhood experiences, like abuse, parental loss, and household dysfunction are 1.6 to 2.4 times more likely to develop chronic diseases, including obesity, cancer, heart disease, stroke, and diabetes, and their life expectancy can be shortened by as much as 20 years.^16, 17^ In a subset of nonhuman primates, including baboons, rhesus macaques, marmosets, and chimpanzees, early-life adversity, including maternal loss, is associated with a shorter lifespan and impaired welfare.^18-22^ Although orphaned elephants have been shown to exhibit altered behavioral responses and impaired access to social partners,^23^ and our own work has demonstrated transiently elevated fecal glucocorticoid metabolite concentrations in the youngest orphans,^24^ it is unclear whether such experiences are associated with accelerated aging or have long-lasting physiological imprints.

In humans, epigenetic pathways have emerged as a key mechanistic link between early-life experiences and later-life health outcomes, offering new insights into how they can leave lasting biological residues and influence disease risk.^25^ Epigenetic modifications, such as DNA methylation (DNAm), enable the genome to respond to environmental cues without altering the underlying DNA sequence, often by regulating chromatin structure and gene accessibility.^26^ DNAm, one of the most widely studied epigenetic processes, involves the addition of methyl groups to CpG sites and plays a crucial role in modulating gene expression.^27^ In addition, genome-wide studies have identified differentially methylated regions associated with early-life experiences, including parental loss, particularly in pathways involved in inflammation and immune regulation.^28, 29^ This biological embedding of adversity through epigenetic mechanisms is thought to contribute to increased risk for chronic diseases and shorter lifespan. Whether this occurs in elephants, who share many key social and life-history characteristics with humans, has yet to be investigated.

Epigenetic clocks, which quantify genome-wide DNAm patterns to estimate biological age, have been extensively studied in humans^30^ as well as mice,^31^ non-human primates,^32^ and dogs.^33^ These measures are indices of DNAm sites that show differential methylation associated with chronological age (first generation clocks, those most available for non-human species, including elephants^34,35^) and/or aging-related health indicators (second/third generation clocks, including GrimAge,^36^ DunedinPace^37^). Most second/third generation clocks are trained on blood-derived immune cells (and, in some cases, saliva), although epithelial cells have also been used. DNAm aging using later generation clocks has been linked to traumatic childhood experiences in humans,^38^ including parental loss,^39^ with effects on later-life health.^40^

In this study, we collected skin samples from wild orphaned and non-orphaned (control) savanna elephants residing in Kafue National Park, Zambia. At sampling, most orphans were under human care but foraged daily in the national park. Using the UniversalClock3Skin, we assessed whether orphans showed accelerated aging and an epigenetic signature compared with wild control elephants. In addition, within a subset, we investigated temporal changes in DNAm age. We hypothesized that orphaned elephants would exhibit accelerated DNAm age and a specific epigenetic signature resulting from early-life trauma. We also hypothesized that a 1-year interval between repeated sampling would be sufficient to see changes in DNAm age within individuals, reflecting differences in the pace of epigenetic aging.

## Results

A total of 32 skin samples were collected from 23 African savanna elephants residing in Kafue National Park, Zambia, including 14 orphaned (8 males, 6 females) and 9 non-orphaned wild (2 males, 7 females) elephants. Among these, six orphaned and three non-orphaned individuals contributed repeated samples. The mean interval between samples was 420 ± 93 days (range: 321 – 574 days) for orphaned elephants and 655 ± 100 days (range: 589 – 770 days) for non-orphaned elephants. In total, 20 samples were obtained from orphaned elephants and 12 from non-orphaned elephants (Table 1). All orphans were rescued while still primarily milk-dependent, at ∼2 years of age (mean ± SD = 444 ± 264 days, 92 – 915 days). At the time of sample collection, chronological age ranged from 4 to 17 years (mean ± SD = 9.5 ± 3.94 years; n = 6 females, 8 males) and from 9 to 35 years (mean ± SD= 16.9 ± 9.11 years; n = 7 females, 2 males) for orphaned and non-orphaned elephants, respectively. Following principal component (PCA) and inter-array correlation analyses on DNAm data, one skin sample was identified as an outlier and removed, leaving 31 skin samples for the primary analysis. Because samples were collected at varying intervals following orphaning (mean ± SD = 8.5 ± 3.7 years, 3.0 – 15.6 years), we tested whether time since orphaning was associated with epigenetic age at the time of sample collection, adjusting for sex and chronological age. No significant association was observed (estimate: 0.067, SE = 0.276, p = 0.811; Figure S1, Table S1).

**Table 1.** Demographic and sample data on the study population.

| Elephant ID | Estimated DOB | Sex | Collection Date (mm/dd/yy) | Orphan Status | Date of Orphaning |
| --- | --- | --- | --- | --- | --- |
| Jordan | 2004 | F | 5/11/22 | No |  |
| Blue | 2000 | F | 5/11/22 | No |  |
| Blue | 2000 | F | 6/19/24 | No |  |
| Sonnet | 1987 | F | 5/11/22 | No |  |
| Katongo | 2013 | F | 5/12/22 | No |  |
| Mupondopondo | 2011 | F | 5/12/22 | No |  |
| Kaleni | 2012 | F | 10/22/22 | No |  |
| Kaleni | 2012 | F | 6/18/24 | No |  |
| Kafue | 1997 | F | 10/22/22 | No |  |
| Bangweulu | 2012 | M | 10/23/22 | No |  |
| Musungwa | 2013 | M | 10/23/22 | No |  |
| Musungwa | 2013 | M | 6/3/24 | No |  |
| Batoka | 4/1/2008 | M | 10/23/22 | Yes | 10/09 |
| Chamilandu | 4/1/2006 | F | 5/22/23 | Yes | 11/07 |
| Kasewe | 12/1/2015 | F | 6/1/24 | Yes | 27/9/16 |
| Kavala | 10/1/2010 | F | 3/29/23 | Yes | 8/11 |
| Ludaka | 5/1/2017 | M | 3/29/23 | Yes | 24/10/19 |
| Ludaka | 5/1/2017 | M | 6/20/24 | Yes | 24/10/19 |
| Mosi | 8/1/2010 | M | 4/28/22 | Yes | 8/12 |
| Mosi | 8/1/2010 | M | 5/23/23 | Yes | 8/12 |
| Muchichili | 7/1/2014 | M | 5/12/22 | Yes | 11/1/16 |
| Musolole | 5/1/2011 | M | 4/28/22 | Yes | 10/11 |
| Musolole | 5/1/2011 | M | 3/29/23 | Yes | 10/11 |
| Nkala | 3/1/2013 | M | 3/29/23 | Yes | 6/13 |
| Nkala | 3/1/2013 | M | 6/20/24 | Yes | 6/13 |
| Olimba | 11/1/2018 | F | 10/5/22 | Yes | 30/10/19 |
| Olimba | 11/1/2018 | F | 5/1/24 | Yes | 30/10/19 |
| Fungulani | 11/1/2017 | F | 8/4/23 | Yes | 15/12/18 |
| Mulisani | 3/1/2015 | M | 8/4/23 | Yes | 25/9/17 |
| Mulisani | 3/1/2015 | M | 6/20/24 | Yes | 25/9/17 |
| Tafika | 11/1/2008 | M | 5/22/23 | Yes | 8/09 |
| Lufutuko | 12/1/2017 | F | 3/29/23 | Yes | 6/12/18 |

### Early life trauma is associated with negative epigenetic age acceleration in elephants

Methylation clocks are powerful biomarkers for estimating age and age acceleration. We tested whether experiencing a traumatic event early in life, i.e., orphaning, in elephants was associated with accelerated aging. Using a linear mixed model to account for chronological age, sex, and repeated measures within individuals, we found that orphaned elephants demonstrated a statistically significant negative DNAm age acceleration compared to controls (estimate = -1.934, SE = 0.729, p = 0.013; Figure 1; Table 2). Because chronological age differed substantially between groups, we conducted sensitivity analyses in an age-matched subset of orphans and non-orphans (age: 9 - 18 years; n = 7 and 6, respectively). Chronological age did not differ between groups in this subset (p = 0.610). Even within this smaller, age-matched sample, the negative age acceleration in orphans compared to non-orphans remained statistically significant (estimate = - 4.359, SE = 1.769, p = 0.039; Figure S2). We further assessed whether age acceleration could reflect nonlinearity or heteroskedasticity in the epigenetic clock by plotting residuals against chronological age (Figure S3). Residuals were distributed on both sides of zero across the age range, suggesting the findings were unlikely to be a mathematical artifact of clock bias.

**Figure 1.**
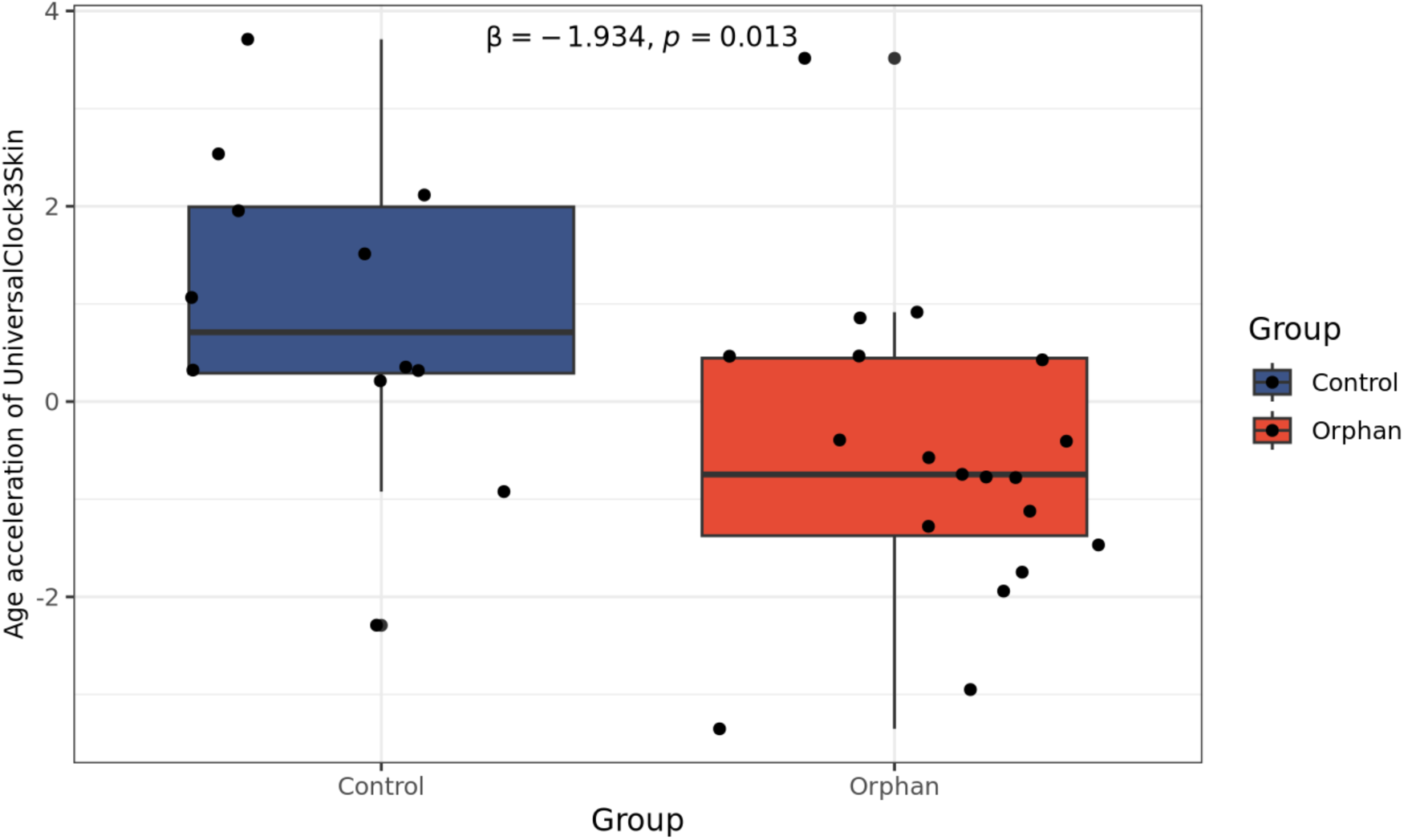
Epigenetic age acceleration in orphaned versus non-orphaned elephants in skin tissue. Age acceleration was calculated as the residual from a linear mixed-effects model of skin DNAm age (UniversalClock3Skin) regressed on chronological age, with Elephant ID included as a random effect to account for repeated measures. Individual data points represent separate sampling events; the boxplot displays the median, interquartile range (IQR), and whiskers extending to 1.5×IQR. The reported p-value and effect size (Estimate) were derived from a subsequent linear mixed-effects model comparing groups while adjusting for age and sex (β=−1.934, p=0.013). Colors denote group status: Blue (Control, n=19) and Red (Orphan, n=12).

**Table 2.** Coefficients of LME model ‘AgeAccel ∼ 1 + Group + Age + Sex + (1 | Elephant). P-values <0.05 are bolded.

|  | Estimate | Std. Error | DF | t-value | p-value |
| --- | --- | --- | --- | --- | --- |
| Intercept | 2.304 | 0.908 | 27 | 2.536 | **0.017** |
| Group | -1.934 | 0.729 | 27 | -2.654 | **0.013** |
| Age | -0.080 | 0.047 | 27 | -1.712 | 0.100 |
| Sex | -0.349 | 0.632 | 27 | -0.552 | 0.586 |

The reference group for “Group” was non-orphans; The reference group for “Sex” was female.

We assessed whether an approximately 1-year time period was sufficient to observe changes in DNAm aging within an individual (n = 6 orphans; n = 3 non-orphans; Table 1) and whether there was a difference in the rate of change in age acceleration between groups. Using a linear mixed-effects model, we found that orphaned elephants exhibited significantly lower baseline age acceleration than controls (estimate = -2.925, SE = 1.254, p = 0.038; Table 3). However, there was no significant change in epigenetic age acceleration between the first and second time points between groups (estimate = -0.171, SE = 1.23, p = 0.892, Figure S4) or when all individuals were combined (estimate = 1.194, SE = 0.793, p = 0.154, Figure S5). The interaction between time and group was not significant (estimate = 2.046, SE = 1.507, p = 0.199; Table 3), indicating that the rate of change in age acceleration did not differ between orphaned and non-orphaned elephants over the study period. Further, initial DNAm age did not significantly influence longitudinal change in age acceleration (estimate = -0.141, SE = 0.094, p = 0.157). There was no evidence of sex differences in longitudinal change in age acceleration (estimate = -0.005, SE = 0.872, p = 1.000).

**Table 3.** Coefficients of LME model ‘AgeAccel ∼ 1 + Timepoint_F * Group + Baseline_Age + Sex + (1 I ElephantID)’. P-values <0.05 are bolded.

|  | Estimate | Std. Error | DF | t-value | p-value |
| --- | --- | --- | --- | --- | --- |
| Intercept | 2.724 | 1.508 | 12 | 1.806 | 0.096 |
| Timepoint_Fsecond | -0.171 | 1.230 | 12 | -0.139 | 0.892 |
| Group | -2.925 | 1.254 | 12 | -2.332 | **0.038** |
| Baseline_Age | -0.141 | 0.094 | 12 | -1.510 | 0.157 |
| Sex | -0.005 | 0.872 | 12 | -0.006 | 0.996 |
| Timepoint_Fsecond:Group | 2.046 | 1.507 | 12 | 1.358 | 0.199 |

Timepoint\_Fsecond: The estimated average change in AgeAccel from the first to the second time point, within the reference group (non-orphans); Group: the estimated difference in baseline AgeAccel (at the first time point) between the orphan and non-orphan group; Baseline\_Age: The expected change in AgeAccel per one-unit increase in Baseline\_Age, holding all other variables constant; Sex: the difference in AgeAccel between male and female elephants (assuming female is the reference sex); Timepoint\_Fsecond:Group: The key term. This is the difference in the change rate (T2-T1) between the orphan and non-orphan group.

### Early life adversity contributes to a small epigenetic signature in elephants

We performed an epigenome-wide association study (EWAS) to determine whether early orphaning leaves a discernible methylomic signature. A small subset of CpG sites (n = 13, q < 0.05, Table S1) demonstrated significant differential methylation between orphaned and non-orphaned elephants, characterized primarily by hypomethylation in orphans (Figure 2A). The p-value distribution for the group comparison showed a distinct peak at very low values but was otherwise approximately uniform (Figure 2B), indicating that while trauma-associated changes are present, they are localized to specific sites rather than representing systemic methylomic shifts. While hypomethylation predominated among the significant sites (Figure 2C), we observed no global directional bias across the epigenome (Fisher’s exact test: p = 0.405, OR = 0.516). Genes that did significantly differ by group included hypomethylation of *POU2F2, AUTS2, PCDHGC5, PIM1*, *GLI4,* and *TRMT1*, and hypermethylation of *LRRC7, PRKG2, EIF4E3*, and *PJA1* (q < 0.05).

**Figure 2.**
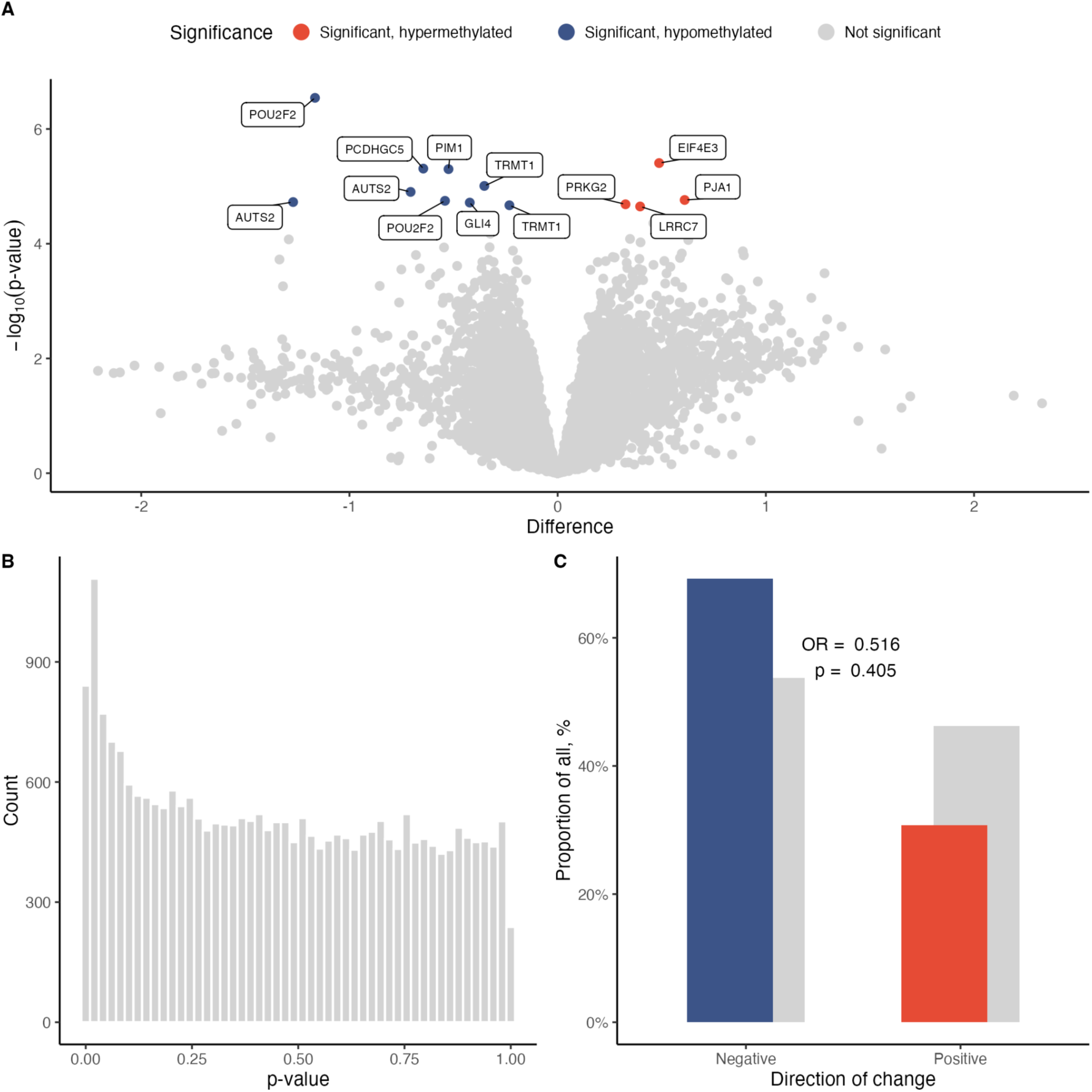
Epigenome-wide differential methylation associated with early-life trauma. (A) Volcano plot illustrating the distribution of CpG-specific methylation differences between orphaned and control elephants. Labelled points denote the 13 significantly differentially methylated CpGs (q<0.05) and their associated gene symbols. Significant hypomethylation is indicated in blue, hypermethylation in red, and non-significant sites in gray. (B) Histogram of p-values derived from the linear regression model. (C) Directional bias of differential methylation. Bar plots represent the proportion of hypermethylated (red) and hypomethylated (blue) sites among significant CpGs versus non-significant background sites (gray). Fisher’s exact test was used to assess the tendency towards hyper- or hypomethylation. The odds ratio (OR) represents the result from Fisher’s exact test.

Secondary analyses tested a total of 25,823 CpGs for association with chronological age. Age was not significantly associated with differential CpG methylation at the genome-wide level after correcting for multiple testing (q > 0.05). There were some age trends at uncorrected p-values (1,447 CpGs were significant at p < 0.05; 25 CpGs were significant at p < 3.16x10^-4^) but the lack of CpGs passing FDR suggests insufficient power to detect age effects at genome-wide significance.

A gene set enrichment analysis (GSEA) based on the DNAm differences between orphaned and non-orphaned elephants was conducted to ascertain which biological changes accompany early life trauma. We identified nine pathways significantly affected in orphaned elephants (Table S2). These primarily included developmental regulation pathways (NES = -2.866; -2.735, q = 0.014; 0.016), which were significantly enriched with hypomethylated genes in orphans. Additional pathways associated with signaling and cellular organization were also enriched with hypomethylated genes (Figure 3). Conversely, positively enriched pathways included negative regulation of leukocyte activation (NES = 2.819, q = 0.015), cell-cell signaling (NES = 3.121, q < 0.001), and hallmark KRAS signaling NES: 3.126, q < 0.001).

**Figure 3.**
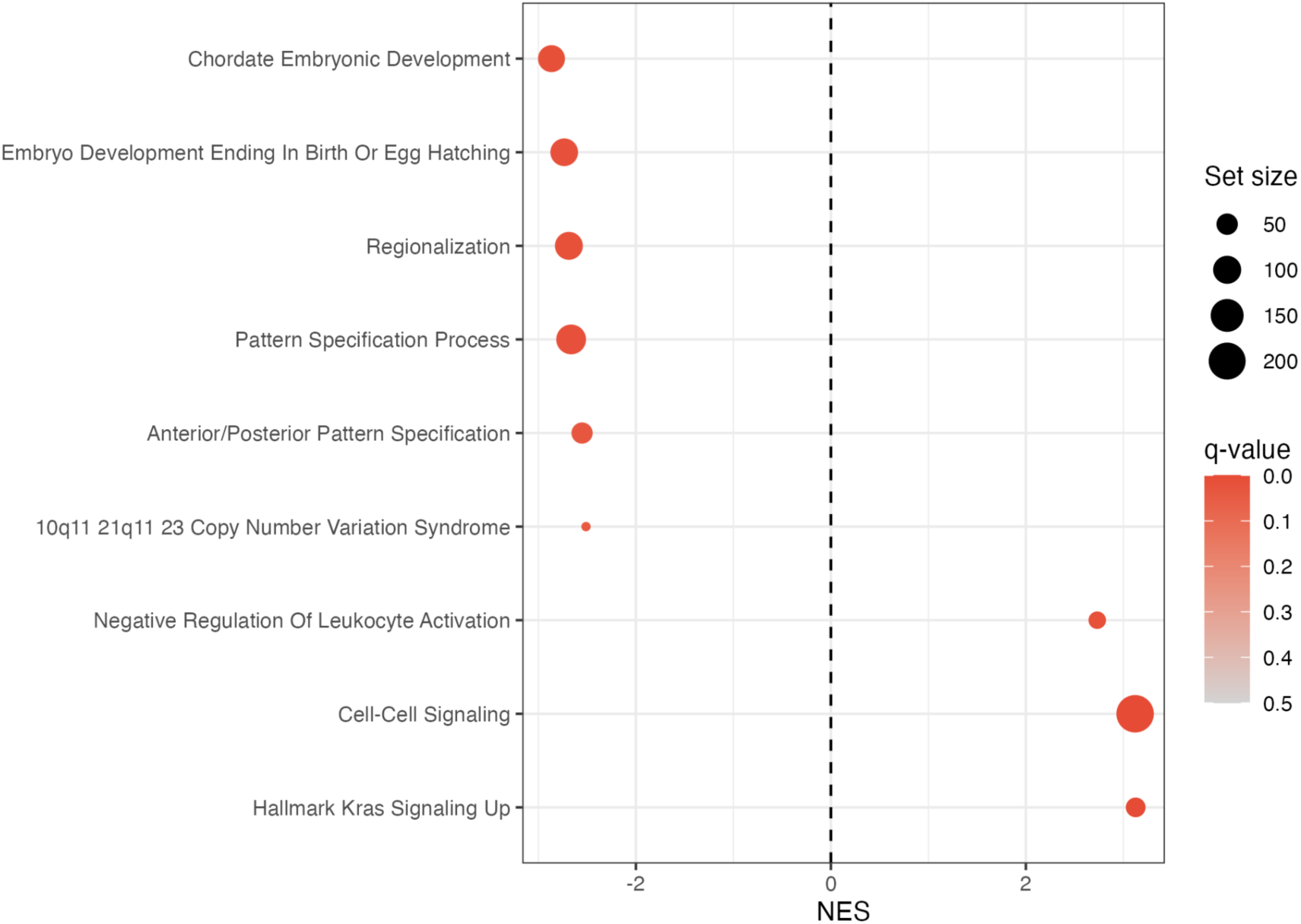
Statistically significant gene set enrichment analysis (GSEA) results. Node size reflects the number of genes identified within each pathway. The bubble plot highlights significantly enriched pathways, with bubble size proportional to pathway size and color indicating statistical significance (*q* < 0.05).

## Discussion

This study examined DNAm age in elephants residing in Kafue National Park, Zambia, that had experienced early-life trauma — maternal loss — and whether epigenetic profiles differed between orphaned and non-orphaned individuals. Orphaned elephants exhibited significantly lower epigenetic age acceleration compared to non-orphaned elephants based on skin samples. Despite this initial divergence, longitudinal analysis confirmed that both groups maintained a similar rate of epigenetic aging over a 1-year interval. Only a few CpG sites differed significantly between groups, with a trend toward hypomethylation in orphans, suggesting a subtle but nonsignificant directional association. Together, these findings suggest that early maternal loss in this population does not accelerate epigenetic aging and may even slow it, while leaving a subtle but targeted epigenetic imprint localized to specific developmental and signaling pathways. This finding was unexpected and ran counter to our initial hypothesis that early-life trauma would hasten biological aging.

Several factors may explain why orphaned elephants showed significantly lower epigenetic age acceleration. First, it is possible that an age bias is driving the observed association, given our small sample size. However, sensitivity analyses in age-matched individuals yielded the same results: orphans exhibited lower epigenetic age acceleration, suggesting the finding is unlikely to be an artifact of the clock itself. Second, and importantly, most orphans were still living under human care at the time of sampling (11 out of 14 individuals). At the rescue facility, orphans had access to a constant water source, received dietary supplementation and veterinarian care as needed, and were accompanied by humans during foraging excursions. These conditions likely reduced their exposure to environmental and nutritional stress compared to non-orphans, whose vital needs were less secure. By contrast, non-orphaned elephants traveled greater distances (unpublished data) to find food and water and were solely responsible for protecting themselves and their calves from predation, reflecting a more energetically demanding lifestyle. Anecdotal reports describe the Kafue elephants as highly stressed, based on responses to humans; however, fecal glucocorticoid metabolite concentrations did not differ from those previously measured in the same orphaned individuals.^24^ It is possible then that differences in perceived stress alone may not fully account for the older DNAm age observed in non-orphaned elephants. A third, non-mutually exclusive explanation is that maternal loss trauma may alter developmental trajectories, resulting in delayed biological maturation rather than accelerated aging. In humans and other mammals, exposure to severe stress or trauma during early life has been associated with the persistence or re-emergence of developmentally earlier behavioral and physiological states, including increased dependency, altered social behavior, and shifts in neuroendocrine regulation.^20, 41-43^ Such patterns have been interpreted as adaptive responses to adverse environments, prioritizing survival over growth and maturation.^44^ In this context, the younger-than-expected epigenetic age observed in orphaned elephants may reflect a delay in biological development rather than a protective slowing of age per se. If so, DNAm age in this system may capture aspects of developmental timing in addition to chronological aging.

Distinguishing between delayed maturation and true deceleration of aging will require longitudinal data spanning key developmental transitions over extended periods. Continued longitudinal sampling of orphans as they transition from human care to fully wild conditions will also clarify whether lower epigenetic age acceleration persists. Another consideration is potential bias within our orphan sample. Field staff estimate that roughly 31% of rescued orphans succumb to physical and emotional injuries within 1 month of rescue. Those that do survive may possess unique resilience traits,^45^ which could contribute to the younger DNAm age at early life stages. Future work should compare DNAm age and epigenetic profiles between surviving and non-surviving orphans to test this hypothesis. Ultimately, something unique to orphaned elephants may underlie the younger DNAm age, a pattern that contrasts with most other mammals.^46-48^

Orphaned elephants showed significantly lower epigenetic age acceleration at the first observation; however, among individuals with two measurements approximately one year apart, we found no evidence that orphaning was associated with differential 1-year change in epigenetic age acceleration. This pattern is consistent with evidence that epigenetic aging measures exhibit moderate-to-high stability across repeated assessments,^49, 50^ while still allowing for environmentally driven variation. In humans, traumatic stress and adverse childhood experiences have been linked to higher epigenetic age acceleration or faster pace measures in many studies,^40^ although meta-analytic findings indicate effect sizes are generally small and heterogeneous across clocks and study designs.^51, 52^ One possible explanation for why the timing of orphaning was not related to DNAm age in this study was that the 1-year sampling interval was too short to capture measurable changes, especially given elephants’ long lifespans. Alternatively, any changes may have been too small to overcome the inherent error in the epigenetic clock due to the small sample size. Although not significant, a larger sample size and longer intervals between repeated measures would allow a more robust assessment of “set-point” differences from changes over time. To address these limitations, we are currently collecting additional samples to increase the sample size and extend the time frame for repeated measures.

Notably, the biological pathways identified in our analysis, particularly those related to developmental regulation, overlap with pathways previously shown to be associated with chronological aging in elephants.^34^ However, in contrast to prior work demonstrating widespread age-associated methylation changes across thousands of CpG sites in captive elephants,^34^ we detected only a small number of differentially methylated sites in our wild population. These sites were primarily associated with developmental and transcriptional pathways, which were significantly enriched with hypomethylated genes in orphans, suggesting potential effects on gene expression linked to growth and development. This pattern could reflect stress-related developmental plasticity, potentially indicating prolonged or altered activation of developmental processes consistent with delayed biological maturation.^52^ Conversely, positively enriched pathways included negative regulation of cell activation, cell-cell signaling, and hallmark KRAS signaling, potentially reflecting suppressed immune responses and altered signaling pathways. Both processes have been linked to later-life health consequences following early trauma. However, given the limited number of differentially methylated sites and the absence of gene expression data, these interpretations should be considered speculative.

This study has several limitations that should be considered when interpreting the results. First, the sample size was modest and included individuals with repeated measures, which may limit statistical power and the ability to detect subtle differences, particularly in genome-wide analyses. Second, orphaned and non-orphaned elephants differed in chronological age, raising the possibility that age-related processes could confound group comparisons. To mitigate this, we used a residual-based age acceleration metric and included chronological age as a covariate in all models, and importantly, sensitivity analyses restricted to an age-matched subset yielded consistent results, suggesting that the observed associations are unlikely to be solely driven by age differences. However, we cannot fully exclude the possibility of non-linear age effects or residual confounding, particularly given the limited sample size. Third, orphaned individuals were sampled at varying time points following the orphaning event, which may have limited our ability to detect transient changes in epigenetic aging. It is possible that early-life trauma induces an initial period of accelerated aging that subsequently stabilizes or normalizes over time, a pattern that would not be captured with cross-sectional sampling at heterogeneous intervals.

Although we tested for an association between time since orphaning and epigenetic age acceleration and found no significant relationship, this analysis may have been underpowered to detect non-linear or time-dependent differences. We found little evidence of a persistent epigenetic residue of early-life experiences across the life course, suggesting that any epigenetic age differences may be transient or below our detection threshold and therefore may have limited implications for long-term health outcomes. Fourth, although we applied established statistical approaches to control for technical and biological variation in the EWAS (including RUVSeq, duplicateCorrelation, and multiple testing correction), the potential for p-value inflation and reduced stability of effect estimates in small samples remains a concern. As such, the identified CpG sites and enriched pathways should be interpreted cautiously and viewed as hypothesis-generating rather than definitive. Relatedly, gene set enrichment results depend on the underlying EWAS signal and may reflect biases inherent to the array design or limited power. Finally, the use of a two-step residual approach to estimate epigenetic age acceleration, while widely used, may complicate interpretation and assumes primarily linear relationships between chronological and epigenetic age. Although diagnostic plots did not indicate systematic bias, more complex modeling approaches and larger datasets will be needed to fully evaluate potential non-linear dynamics. Future studies with larger sample sizes, broader age overlap, and extended longitudinal follow-up will be essential to confirm these findings and further elucidate the epigenetic consequences of early-life trauma in elephants.

In sum, our data suggest that orphaned elephants do not exhibit accelerated aging or DNAm age older than expected compared to non-orphaned elephants living in the same national park. This pattern contrasts with patterns observed in most other species and may reflect unique biological adaptations that buffer against epigenetic consequences of early maternal loss. Given their longevity, shared life-history traits with humans, and distinctive biology, elephants offer an invaluable comparative species for identifying mechanisms of resilience and vulnerability, with potential relevance to human health.

## Materials and Methods

### Ethics statement

This study was approved by the Indiana University Animal Use and Care Committee (22-022) and Zambia’s Department of National Parks and Wildlife (DNPW; NPW/8/27/1). Samples were exported under Zambia’s CITES export permit (21331) and imported under the United States of America’s CITES import permit (23US39890E/9).

### Animals and setting

This study was conducted in collaboration with Game Rangers International (GRI), Zambia’s DNPW, and African Parks (AP). GRI is a conservation organization that works with DNPW, and through their Elephant Orphanage Project, they rescue and rehabilitate orphaned elephant calves, and ultimately release them back into the wild in Kafue National Park, Zambia. We studied 23 African savanna elephants residing in Kafue National Park, Zambia: 14 orphaned (8 males, 6 females) and 9 non-orphaned wild (2 males, 7 females) elephants. At the time of rescue, all orphans were less than 24 months old, with age estimated to within 3 months based on key developmental features.^53^ At approximately 3-4 years of age, orphans are translocated from the nursery to the Kafue Release Facility (KRF), which is located within Kafue. Ultimately, orphaned elephants are outfitted with GPS collars to allowing tracking once they leave the KRF. Orphan demographics are outlined in Table 1. In May and October of 2022, non-orphaned wild elephants residing within 20 km of the KRF were anesthetized and outfitted with GPS collars (Table 1). Age was estimated based on shoulder height,^48^ measured at time of GPS collaring.

### Sample collection

Between May 2022 and June 2024, when elephants were anesthetized for either translocation or GPS collaring, a small piece of skin tissue was cut from the elephant’s ear and immediately placed in DNAShield (Zymo Research, product #R1100-250). Vials were labelled with elephant ID, date and time of collection, and then stored at -20° C until shipment on ice to the United States for DNA extraction.

In addition to skin samples collected when an elephant was anesthetized, focal elephants were targeted for skin tissue biopsy darting (Motsumi 5cc biopsy marking dart 1 ½” or 1 ¼” C type, Pretoria, South Africa). Approximately 4x40 mm skin punch biopsy was collected from the elephants’ rear by a Zambia DNPW veterinarian (Figure S6). Darting occurred approximately 50 m from the target animal by vehicle. The biopsy dart released a purple mark at the site of entry, immediately identifying the sampled elephant. Once the elephant started moving, the biopsy dart fell out and was recovered. In the field, the skin sample was immediately placed in a labelled vial with DNAShield and stored as outlined above for skin samples.

### DNA extraction

Genomic DNA was extracted using the Quick-DNA HMW MagBead Kit (Zymo Research, Irvine, CA, USA) according to the manufacturer’s protocol.

### DNAm profiling and quality control

Genome-wide DNA methylation was measured from 250 ng of bisulfite-converted genomic DNA using the custom Illumina “HorvathMammalMethylChip40” array, which provides high coverage of approximately 36,000 highly conserved CpG sites across mammalian species and skin tissue.^54^ Raw intensity data were normalized using the SeSAMe R^55^ package to generate methylation estimates (beta values) and detection p-values for each probe. To minimize technical variance and batch effects, repeated samples from the same individual samples were processed on the same array chip. Rigorous quality control was performed to ensure data integrity. Probes were marked as failed if both methylated and unmethylated channels reported background signal levels (p > 0.01). Samples with a scaled proportion of failed probes greater than 2 were excluded as outliers. For downstream epigenome-wide association studies (EWAS), probes failing in more than 75% of samples were removed. Further outlier detection was conducted via Principal Component Analysis (PCA) on the centered, unnormalized beta-value matrix; samples exceeding 2 standard deviations (SD) on the first three principal components were excluded. Finally, hierarchical clustering based on correlation distance was used to identify and remove any remaining samples situated more than 2 SD from the mean. After QC, 31 skin samples remained for analysis.

### Measure of epigenetic age estimation and acceleration

Epigenetic age was estimated using the UniversalClock3Skin for skin samples.^56^ To determine whether there was a difference in DNAm age by group status, age acceleration (AgeAccel) was calculated for each skin sample using a two-step residual method, performed only on individuals with repeated measures (i.e., multiple samples). Age was included as a covariate in the second-step model to specifically account for any remaining linear relationship between age and the residuals. Within the retained dataset, samples for each individual were ordered by chronological age to define the time points: Timepoint_F: a factor variable created to distinguish the collection time, coded as first and second (for the earliest and next sample, respectively). A continuous covariate, the individual’s age at the first time point, was created and termed Baseline_Age. After filtering, a linear mixed effects (LME) model was fit to define the expected relationship between the UniversalClock3Skin’s output and chronological age. This baseline model induced age as a fixed effect and a random intercept for elephant ID to account for non-independence of samples from the same individual. This model was fit using Maximum Likelihood (ML) estimation.

Second, the AgeAccel variable was defined as the residuals from this baseline model. A positive residual indicated an DNAm age older than expected (‘positive age acceleration’), while a negative residual indicated an age younger than expected. Lastly, this AgeAccel was used as the outcome variable in subsequent LME models, fit with Restricted Maximum Likelihood (REML) estimation to test for a differential change in age acceleration between the two groups, AgeAccel ∼ 1 + Group + Age + Sex + (1 | ElephantID) and AgeAccel ∼ 1 + Timepoint_F*Group + Baseline_Age + Sex (1 | ElephantID). Secondary analyses included running the same model in a subset of age-matched orphaned and non-orphaned elephants. A t-test was used to ensure groups were age-matched. We also, investigated whether time since orphaning associated with age acceleration, AgeAccel ∼ 1 + TimeSinceOrphaning/365 + Sex + Age + (1 | ElephantID). To assess whether there were any differences between time point 1 and time point 2, regardless of orphan status, we employed the model AgeAccel ∼ 1 + Timepoint_F + Baseline_Age + Sex + (1 | ElephantID). All analyses were performed in R using the lme4^57^ package.

We evaluated model assumptions by examining residual plots of AgeAccel by chronological age for nonlinearity and heteroskedasticity.

### EWAS of orphan status

To identify probes that have significantly different methylation between groups, we used linear modeling. Normalized probe methylation values were used to fit the linear regression model using R package limma.^58^ To account for technical variance and unobserved batch effects, we implemented RUVSeq without over-correcting the biological signal.^59^ A subset of probes least likely to be associated with the condition of interest was utilized as a control set to estimate vectors of unwanted variability. We specifically incorporated two RUVg vectors as covariates in the subsequent linear models to effectively adjust for these hidden sources of variation without over-correcting the biological signal. Given the longitudinal nature of our dataset, we utilized the *duplicateCorrelation* function to account for the intra-subject correlation structure among repeated measures. This approach treats individual identity as a random effect, thereby improving the precision of effect estimates. The linear models were fitted using the least-squares methods by running the *lmFit* function. This method minimized the sum of squared differences between observed values and model-predicted values, providing estimates of the model parameters. Empirical Bayes statistics were estimated using the *eBayes* function. The Benjamini-Hochberg (BH) method was used to correct for multiple testing. Correction for multiple testing was needed to control the FDR when conducting multiple statistical tests simultaneously. Probe methylation differences with q-values < 0.05 were deemed significant.

Specific genes were identified using the official *Loxodonta africana* annotation file for the “HorvathMammalMethylChip40” array provided by the Mammalian Methylation Consortium, which maps the specific CpG probes to their corresponding elephant gene symbols (source: https://github.com/shorvath/MammalianMethylationConsortium/tree/main/Annotations%2C%20Amin%20Haghani/Mammals).

### Gene set enrichment analysis (GSEA)

GSEA was performed to identify biological pathways associated with differential DNA methylation. Input data consisted of probe-level results from the EWAS, including log fold-change (logFC) and p-values. To ensure data integrity, only probes mapped to UCSC reference genes were retained. For genes represented by multiple probes, the most significant probe (smallest p-value) was selected to represent the gene, ensuring each gene contributed a single value to the analysis. To utilize established human pathway databases, *Loxodonta africana* gene symbols were mapped to human orthologs using the biomaRt^60^ R package via Ensembl’s BioMart interface. Extracted human Ensembl and HGNC gene names were subsequently converted to Entrez Gene IDs using the org.Hs.eg.db annotation database for compatibility with downstream tools. Each gene was assigned a ranking metric (RankScore) to incorporate both the direction and statistical significance of methylation changes:

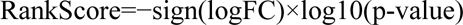

In this formulation, genes with low p-values and increased methylation (positive logFC) are positioned at the top of the ranked list, while those with low p-values and decreased methylation (negative logFC) are at the bottom. Enrichment was calculated using GSEA software.^61, 62^ The analysis was restricted to pathways containing between 10 and 500 genes. The algorithm was executed using the "classic" scoring scheme and "meandiv" normalization method with 1,000 permutations, which reshuffles the data to determine how likely it is to get these results by chance. This acted as a primary control to ensure the results were robust and not random noise. Multiple testing was accounted for using the Benjamini-Hochberg (BH) correction, with significance defined as an q<0.05. A positive Normalized Enrichment Score (NES) indicates a trend toward hypermethylation within a pathway, while a negative NES indicates predominant hypomethylation.

## Data availability

Data will be made publicly available upon publication of the manuscript.

## Acknowledgments and funding

We graciously thank Zambia’s Department of National Parks and Wildlife, African Parks, and Game Rangers International staff and keepers for their support and assistance with making this study possible. We also want to thank former research assistants Constance Banda, Mary Muyoyeta, and Vincent Abere. DEC was supported in part by a National Institute on Aging Award (K01AG072615), the Animal Models for the Social Dimensions of Health and Aging Research Network (NIH R24AG065172), and Indiana Clinical and Translational Sciences Institute grants (UL1TR002529; UM1TR004402).

