## Supplemental File for "Epigenetic Resilience to Early-Life Maternal Loss in African Savanna Elephants"

Table S1. Coefficients of LME model ‘AgeAccel ~ 1 + TimeSinceOrphaning/365 + Age + Sex + (1 | Elephant). P-values <0.05 are bolded.

| | Estimate | Std. Error | DF | t-value | p-value |
|---|---|---|---|---|---|
| Intercept | -0.317 | 1.068 | 15 | -0.297 | 0.770 |
| Time Since Orphaning | 0.067 | 0.276 | 15 | 0.243 | 0.811 |
| Age | -0.090 | 0.270 | 15 | -0.333 | 0.744 |
| Sex | -0.038 | 0.976 | 15 | -0.038 | 0.970 |

The reference group for “Sex” was female.

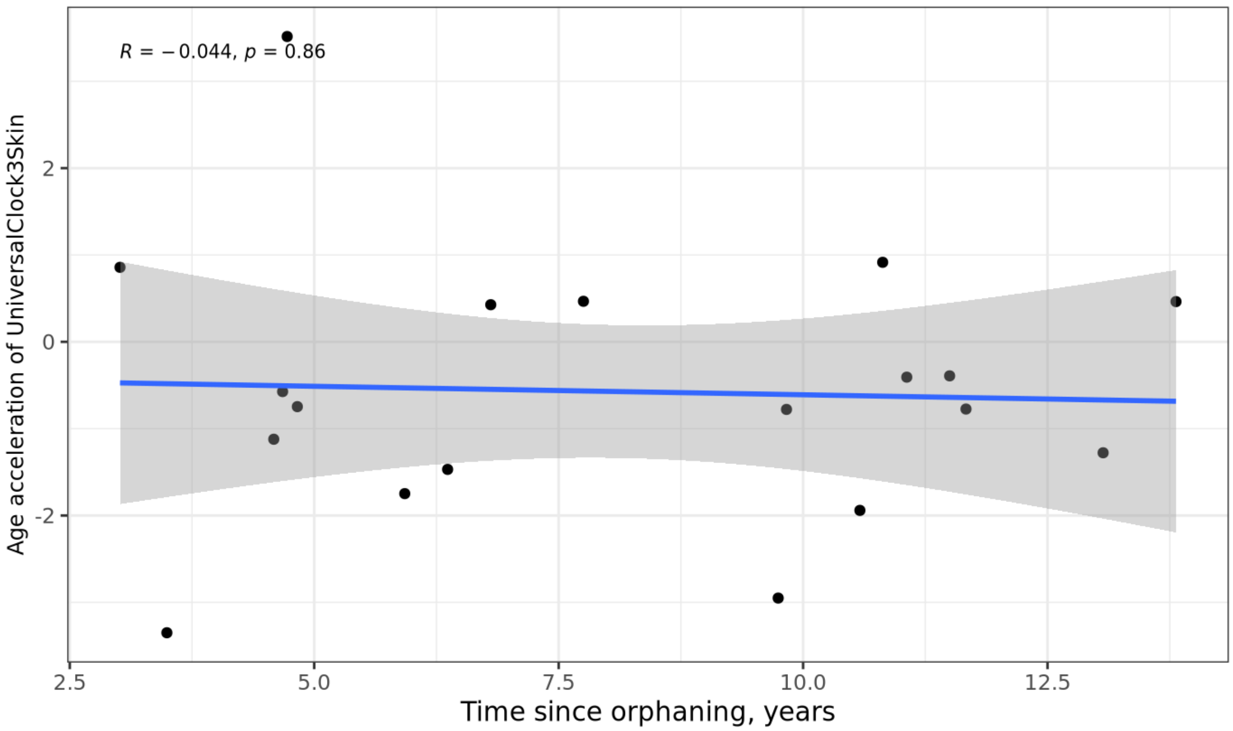


**Figure S1. Time since orphaning and age acceleration in orphaned elephants based on skin samples.** The date of the orphaning event was known for each individual and time since orphaning was calculated as the difference between date of sample collection and date of the orphaning event.

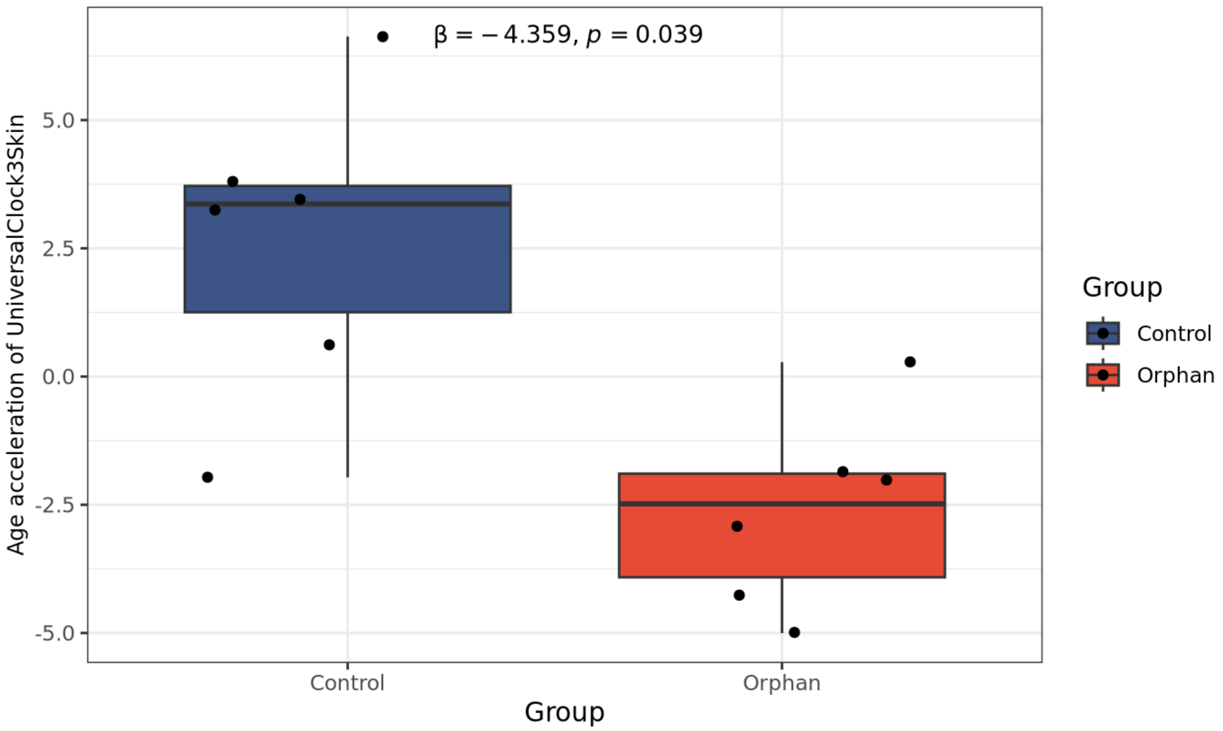


**Figure S2. Epigenetic age acceleration in subset of age-matched orphaned versus non-orphaned elephants in skin tissue.** Chronological age ranged between 9 and 18 years of age. Age acceleration was calculated as the residual from a linear mixed-effects model of skin DNAm age (UniversalClock3Skin) regressed on chronological age. Individual data points represent separate sampling events; the boxplot displays the median, interquartile range (IQR), and whiskers extending to 1.5×IQR. The reported p-value and effect size (Estimate) were derived from a subsequent linear mixed-effects model comparing groups while adjusting for age and sex ($\beta = -4.359$, $p = 0.039$). Colors denote group status: Blue (Control, n=6) and Red (Orphan, n=7).

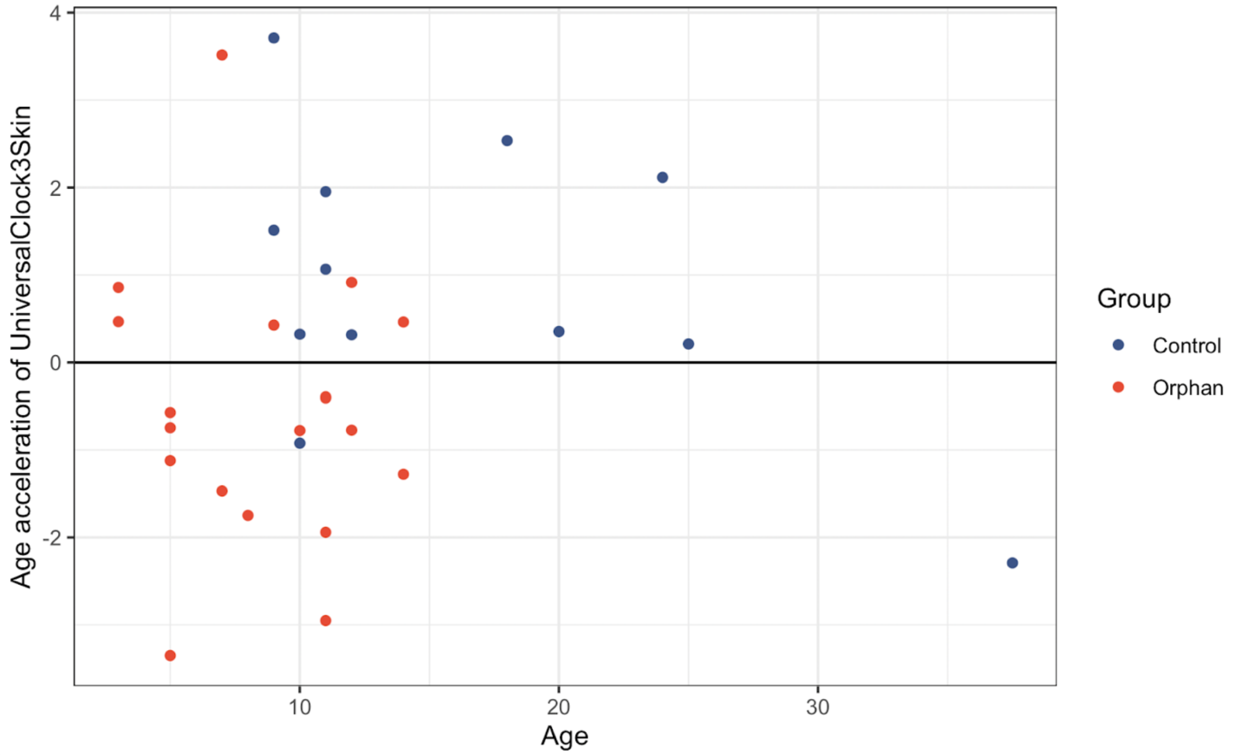


**Figure S3. AgeAccel residuals by chronological age in years for oprhaned and non-orphaned elephants.** Each point represents a skin sample, with blue indicating non-orphaned (control) elephants and red indicating orphaned elephants. The horizontal line at zero represents no age acceleration (i.e., epigenetic age equal to chronological age).

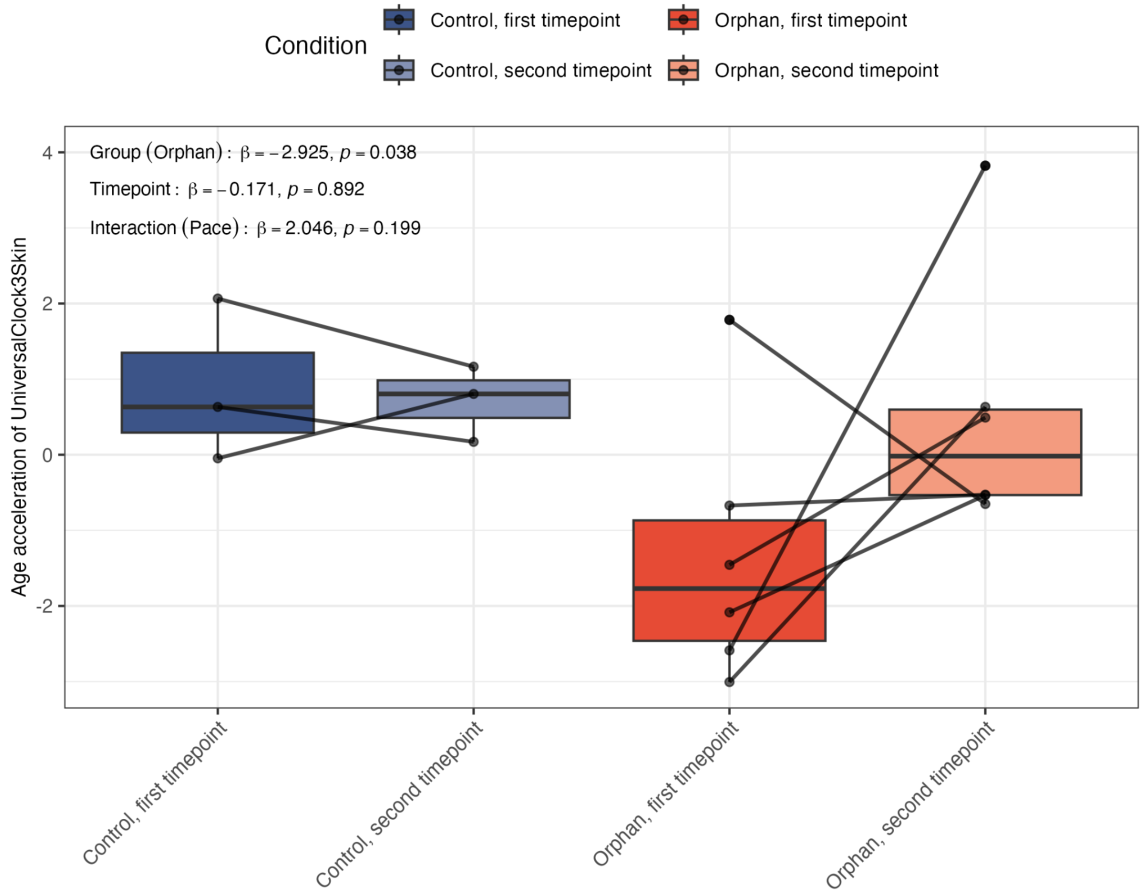


**Figure S4. Longitudinal stability of the epigenetic trauma signature.** Comparison of epigenetic age acceleration in orphaned and control elephants across two timepoints (one-year interval). Statistical annotations (β and p) represent the effect of group status, sampling timepoint, and their interaction, derived from a linear mixed-effects model adjusting for sex and baseline chronological age with Elephant ID as a random effect. Individual trajectories are shown with connecting lines to highlight intra-subject stability. Boxplots represent the median and IQR for each group and timepoint. Colors denote group and sampling period.

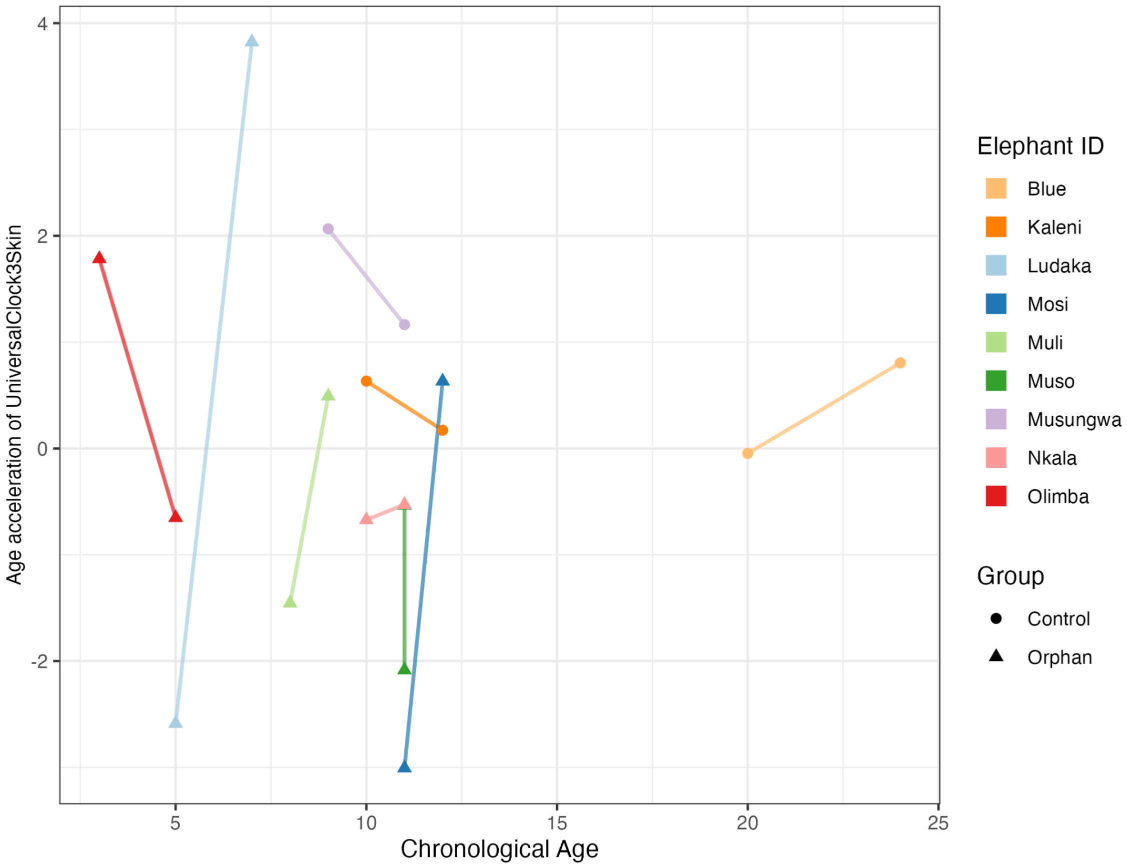


**Figure S5. Within individual change in epigenetic age by chronological age.** Epigenetic age acceleration in orphaned and non-orphaned African savanna elephants across chronological age. Points represent individual elephants, with circles indicating non-orphaned (control) individuals and triangles indicating orphaned individuals. Colors denote elephant ID. Lines connect repeated measures from the same individual across sampling timepoints. Age acceleration was calculated as the residual from the UniversalClock3Skin regressed on chronological age.

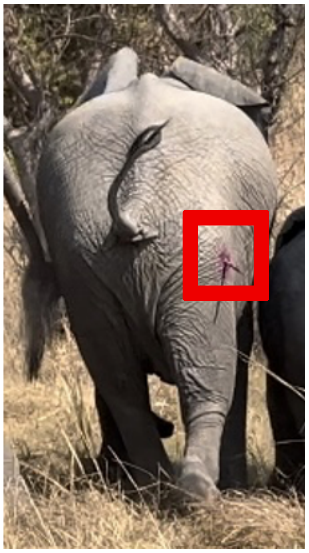

**Figure S6. Example of a dart biopsy of an elephant.** The purple paint provides indication that the elephant was hit with the dart. Once the elephant starts moving, the dart falls out, and then is recovered by the team. Red square highlights the dart.
